# Barium titanate piezoelectric nanoparticles induce M1 polarization in mouse macrophages via ultrasound *in vitro*

**DOI:** 10.1101/2025.10.02.680098

**Authors:** Timothy Connolly, C. Johnson, A. Chen, K. Kempa, D. Hatt, Michael J. Naughton

## Abstract

Macrophages are critical for the maintenance of immune system homeostasis. They differentiate into distinct functional populations, from pro-to anti-inflammatory phenotype, exhibiting remarkable biological plasticity and responding to both chemical and physical cues to achieve these phenotypes. Controlling macrophage cell phenotypes *in vivo*, with temporal and spatial control, could have significant impact on a wide range of human diseases and ailments associated with inflammation, which range from rheumatoid arthritis and Alzheimer’s to cancer tumorigenesis. Piezoelectrics, materials in which pressure causes a voltage and *vice versa*, represent a potential platform for non-invasive and remote modulation of cells and tissues and, in particular, control of immune cell activation. Here, it is demonstrated that RAW264.7 mouse macrophage cells that have taken up piezoelectric nanoparticles (pzNPs) specifically adopt an M1 cellular phenotype and requisite calcium ion influx upon ultrasound stimulation. One can further identify which cells have taken up pzNPs and which cells adopt an M1 polarization in mixed populations of pzNP-loaded and -unloaded cells. The overall goal is to leverage this novel cellular assay to help improve understanding of how biological cells respond to bioelectric stimulation.

## 1. Introduction

Macrophages are a heterogeneous group of myeloid cells ubiquitously found in all mammalian tissue, where they perform diverse tasks depending on their location and gene expression profile, and are involved in initiating and terminating inflammation. They are mobile white blood cells of the immune system, functioning as terminally-differentiated cells of the mononuclear phagocyte system that also encompasses dendritic cells, circulating blood monocytes, and committed myeloid progenitor cells in bone marrow. Highly phagocytic and capable of ingesting cellular debris and foreign substances, macrophages are versatile cells that can undergo differentiation (phenotypically and functionally) in response to their microenvironment. Subpopulations with different functions range from the so-called M1 type that is pro-inflammatory and anti-microbial to the M2 type that is anti-inflammatory.^[1, 2]^ Monocytes from blood, bone marrow or spleen (M0 type) can be differentiated biochemically into macrophages *in vitro* ^[3,4]^ as well as naturally *in vivo*. In response to different environmental stimuli, macrophages can thus adopt either a heal/growth-promoting (M2) or an antitumor, cell-killing phenotype (M1), and are critical for tissue homeostasis. ^[3]^

These major macrophage types are characterized by expression of cell surface markers, secreted cytokines and chemokines, and transcription and epigenetic pathways. Importantly, the M1↔M2 macrophage “polarization” balance governs the fate of tissues and organs in inflammation or injury. A strategy to modulate / control macrophage polarization (*i.e.*, directing M0 monocytes into M1 or M2, M1 into M2, *etc*.) could be a promising therapeutic modality in several infectious and human inflammatory diseases.^[5,6,7]^ M1 polarization is characterized by the expression of inducible nitric oxide synthase (iNOS),^[8]^ reactive oxygen species, and the cytokine interleukin-12 (IL-12). Conversely, M2 macrophages could inhibit T-cell-mediated anti-tumor immune response, and promote tumor angiogenesis, progression and metastasis. ^[6]^ M1 polarization is typically driven by granulocyte macrophage colony-stimulating factor (GM-CSF), lipopolysaccharide (LPS) and interferon-γ (IFN-γ), while IL-4 and IL-13 induce M2 polarization.^[4,9]^

Recent advances have unraveled the significance of nanoparticle (NP)-based electrical stimulation as an attractive approach to modulate immune cell phenotype and activity.^[10,11]^ Piezoelectric (pz) materials,^[12]^ in which pressure elicits a voltage and *vice versa*, represent a potentially useful NP platform for the non- (or at least minimally-) invasive and remote delivery of local electric cues to cells and tissues. pz materials are characterized by a unique charge polarization by either mechanical stimulation or an electric field.^[11,13,14]^ When an internalized pzNP is subjected to mechanical pressure, it produces an electrical field which can, at a minimum, directly interact with the charged cell membrane, potentially triggering a change in resting potential via modulation of intracellular calcium level, and influencing signaling pathways and responses to stimuli, particularly in relation to inflammation and tissue repair for immune cells.

In this context, prior work has shown that charges released on the surface of piezoelectric β*-*PVDF (polyvinylidene fluoride) membranes induced by ultrasound (uS) irradiation can enhance M1 polarization of macrophages cultured on the membrane.^[^ ^15^ ^]^ Other research demonstrated that pz stimulation of barium titanate-coated Ti_6_Al_4_V surfaces promoted an anti-inflammatory M2 polarization of macrophages and bone repair.^[16]^

Macrophage polarization is routinely regulated by biological / biochemical stimulation in the body, but investigations of the regulatory effect of *physical* stimulation are limited. It is becoming increasingly clear that macrophages have exhibited remarkable plasticity to both chemical and physical cues; however, the exact mechanism(s) and the resulting changes are not well understood. Moreover, the use of pzNPs to modulate macrophages, the topic of interest here, has not been reported. These pzNPs can offer a promising platform for delivering electric signals to cells and tissue in a remote manner, and studying the mechanism of bioelectric macrophage polarization.^[^ ^17, 18, 19]^ We report here the ability to influence macrophage polarization with localized electrical signals on pzNPs internalized by cells and activated by *ex situ* ultrasound, potentially advancing applications in targeting inflammation, cancer treatment, and tissue regeneration. Note that this intracell activation process is anticipated to differ significantly from most other reported bioelectric strategies, which employ extracellular static, pulsed, or oscillating electric fields^[20,21, 22]^ generated by external or implanted electrodes, none of which can provide a local, intracell electrical perturbation as describe herein.

## 2. Experimental

### 2.1 Barium titanate nanoparticle preparation

Barium titanate (BaTiO_3_, hereafter BT) pzNPs show great potential for use in manipulating electrostatic and biophysical properties of cells to induce changes in cell fate.^[23]^ However, significant challenges exist in regard to the NPs’ size distribution and the degree of agglomeration or clumping that limits the reproducible effects on biological cells.^[24]^ It is thus important to develop reproducible methods that allow for the stabilization of single homogeneous dispersions of NP preparations that enable cellular uptake and the ability to detect NPs upon loading into cells.

BT NPs used in this work started with 99.9% BT powder comprising aggregated clusters of nominal 300 nm-diameter NPs (US Research Nanomaterials, Inc., Houston, TX). Average NP size (*n* = 20) has been measured by SEM to be 350 ± 50 nm (the supplier quotes 300 nm via SEM), with a hydrodynamic size determined by dynamic light scattering (DLS) to be ∼346 ± 15 nm (*n* = 75), with polydispersity index of 0.253, while confirmation of the tetragonal crystal (*i.e.* piezoelectric) form was provided via X-ray diffraction data from the supplier. Dispersion of the nanopowder into individual NPs was necessary for efficient cellular uptake and to limit toxicity. BT NP systems have been investigated and it was shown that aqueous suspensions can be stabilized by electrostatic and/or steric mechanisms.^[25,26]^ Polyelectrolyte species, which adsorb onto the surface of the NPs, have been found to promote dispersion.^[27]^ Colloidal stabilization of BT suspensions using polyacrylic acid (PAANH_4_) and of polymethacrylic acid (PMAANH_4_) has also been investigated.^[^ ^28^ ^]^ Hu, *et al*. reported on BT aqueous suspensions using PVA-b-COOH coating,^[5]^ while Li, *et al.* used a solution of polyethylene glycol (PEG) to prevent aggregation while maintaining pz properties.^[19]^ NP clusters must be declumped into single NPs and sterilized for subsequent use in cell culture experiments. Clumped NP aggregates have been reported to be toxic to cells and reduce cell survival rate.^[19,23,24]^ We envisioned that coating the pzNPs in PEG-biotin could allow for multiple beneficial effects in investigations examining pz control of cell phenotypes. First, the PEG molecule, by adhering to the surface of the NP, prevents reclumping due to static forces. Second, it increases the rate of uptake by cells, as cellular uptake of NPs is directly dependent on PEG density.^[28]^ By leveraging a PEG-biotin molecule, it is possible to visualize NP uptake by fluorescent imaging using the streptavidin-conjugated fluorescent molecule streptavidin-fluorescein isothiocyanate (SA-FITC), see **Figure 1(A)**.

**Figure 1.**
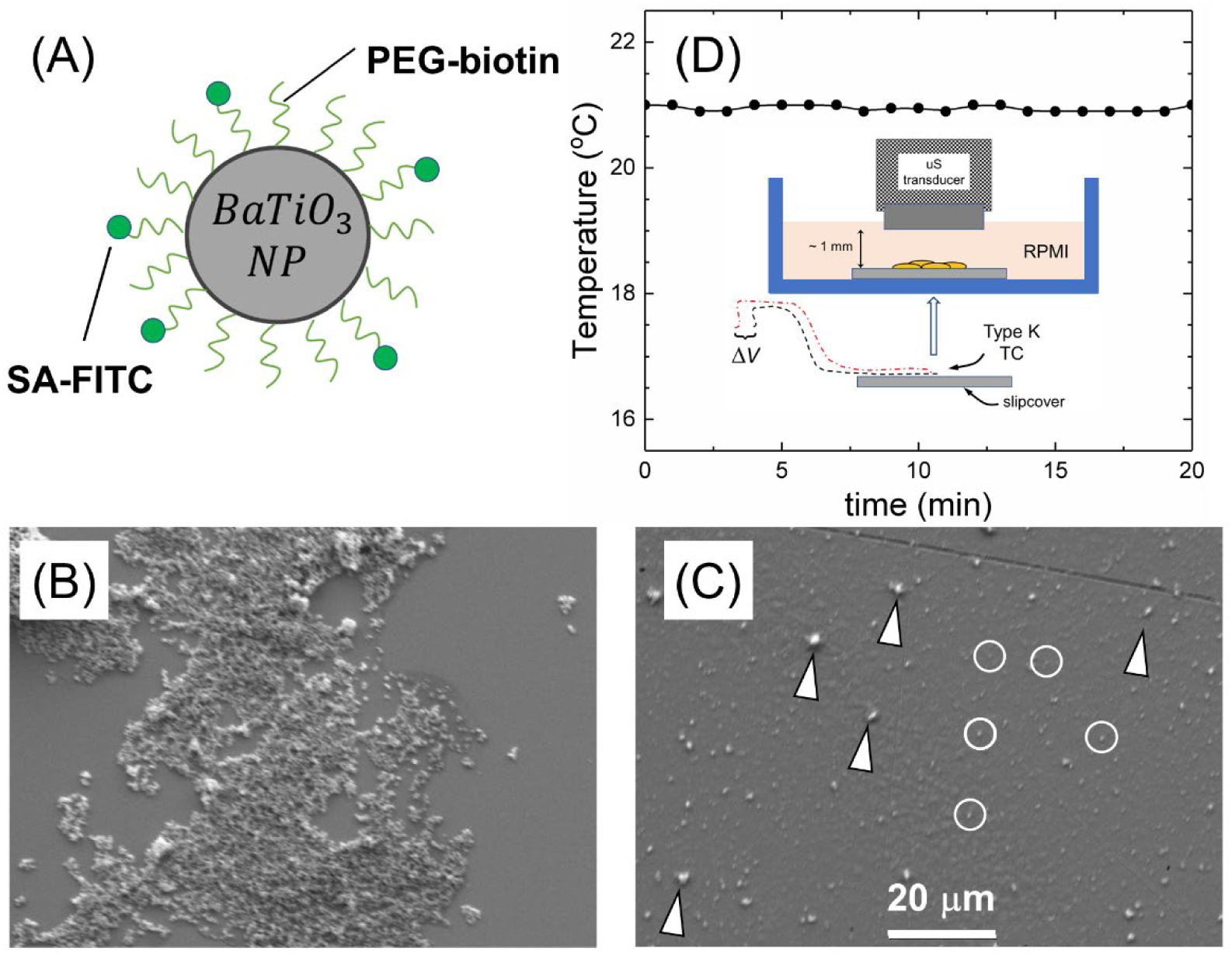
**(A)** Diagram of PEG-biotin coating a barium titanate nanoparticle (NP) showing SA-FITC labeling to PEG-biotin. (**B**) SEM image of untreated, clumped NPs and **(C**) SEM image of declumped NPs after sonication, coated with PEG-biotin and purified prior to cellular uptake studies. Arrows identify representative NP aggregates/clusters, with circles for representative single NPs. (**D**) Results of temperature versus time during application of ultrasound (uS) as measured by a thermocouple (TC) positioned onto a coverslip, immersed in RPMI media, to mimic the experimental conditions of uS simulation of cells. Inset: Schematic of experimental setup of cells on coverslip, and TC layout prior to immersion in media.

We undertook a series of experimental procedures to obtain a solution of sterile individual pzNPs prior to exploring whether these particles would induce pz effects in macrophage cells. For this purpose, we weighed 50 mg of pzNPs from dry powder into a 1.5 ml sterile Eppendorf tube and then resuspended the material in 100 % isopropyl alcohol for 2 h in order to sterilize the particles. All subsequent steps and manipulations were performed with sterile solutions and under a sterile filter hood during solution/media changes. The particles were microcentrifuged at 14,000 rpm (corresponding to ∼17,000 relative centrifugal *g-*force) for 5 min to pelletize the material and then resuspended in a sterile solution of 0.25 M NaCl and 0.005 M PEG-biotin (biotin-PEG-biotin, MW 2,000, BroadPharm Inc., San Diego, CA, no. BP-28703). The suspension of particles then was subjected to sonication using an ultrasonic cleaner for 1 h at room temperature.

After sonication, the pzNPs were washed twice in RPMI media (Roswell Park Memorial Institute 1640 Medium, Gibco Glutamax, ThermoFisher, Inc., No. 61870036) in a microcentrifuge at 14,000 rpm for 5 min and then resuspended in 0.5 ml of sterile RPMI / 10 % - fetal bovine serum media (FBS; Gibco Premium Plus, ThermoFisher, Inc., no. A5669701). We noticed an increased percentage of single pzNPs in the solution. To further optimize the purification protocol, we centrifuged the pzNP solution to a low speed spin of 1,000 rpm (∼1,000 *g*) for an additional 5 min. During this additional centrifugation, we observed that partially clumped pzNPs were pelleted while the single pzNPs remained suspended in solution. We removed the top 250 μl of pzNP solution in RPMI / 10 % FBS to obtain a sterile, purified solution of single NPs that were used for subsequent SEM imaging analysis and cellular uptake studies. SEM imaging was used to characterize the NPs, which clearly show clumped granules of starting powder (Figure 1B) while, after purification as described, a solution containing a majority of single NPs in a sterile solution. Like others, we found that clumped NP aggregates were toxic to cells in culture; however, coating the nanoparticles in PEG prevented reclumping, as NP preparations remained single size and stable for well over a year in solution (as determined by DLS after re-agitation in an ultrasonic bath) (Figure 1C). We calculated a NP concentration to be added to the cells of 20 ng/ml. We next investigated the cellular uptake of these NPs in cell culture.

### 2.2 Cell culture

RAW264.7 cells^[29]^ are a commonly-used mouse macrophage cell line that can be polarized into different phenotypes, including the pro-inflammatory M1 phenotype by stimulation with LPS and IFN-γ. Piezoelectric materials can also influence the polarization of RAW264.7 cells.^[16, 30,31]^ A RAW264.7 cell line was obtained from ATCC (Manassas, VA, no. TIB-71). The cells were thawed from a frozen aliquot and then cultured in T-25 culture flasks containing 5 ml RPMI / 10% FBS media, replaced every 48 h. For uS and experimental manipulations and imaging, the cells were trypsinized and then plated on pre-sterilized glass coverslips (Nunc Thermomax, ThermoFisher, no. 150067) in Corning cell culture dishes at a seeding density of ∼10^5^ cm^−2^. We note that such mild trypsinization has been a published method of passing and culturing cells.^[32,33]^ Cover slips were sterilized in ethanol for 24 h, washed in 1 ml RPMI media prior to being placed in a culture dish. The cells were then allowed to attach to the coverslip and tissue culture plate for 24 h prior to the addition of pzNP nanoparticles and uS treatment.

### 2.3 Ultrasound treatment

Cells were subjected to two rounds of uS stimulation (1 MHz CW frequency, *I*_SATA_ = 0.1 W cm^−2^ power density) of 20 min each, 24 h apart using a Sonitron GTS2000 uS source with a GTSPT06 6 mm-diameter tip transducer (NepaGene Inc., Chiba, Japan via Bulldog Bio, Inc. Portsmouth, NH, USA). Cells were plated on coverslips and placed within cell culture dishes containing 1 ml of RPMI media. An uS probe, sterilized with ethanol and 12 h UV exposure inside the sterile hood, was placed into the culture dish until the probe touched the medium, yielding a measured average probe-cell separation of 0.89 ± 0.06 mm (*n*=20). This probe remained sterile for at least 96 h after it was used within each experimental run. Pressure mapping measurements revealed that the uS intensity at the location of the plated cells was >96% of that at the source and, as shown in Figure 1(D), thermocouple-based temperature-monitoring control experiments (Type K thermocouple, Fluke Model 52 meter) showed no detectable change in temperature (Δ*T* < 0.1 °C) during 20 min stimulation (see also schematic in Figure 1(D) inset). Ultrasound stimulation experiments were performed in a sterile tissue culture hood at room temperature. While a detailed examination of the minimum uS stimulation time required to affect measurable change in cell polarization was not performed here, the aforementioned two rounds were chosen based on prior work that similarly employed multiple rounds.^[15,48]^ Also shown in the inset to Figure 1(D) is a schematic of the configuration of uS application to cells under study.

### 2.4 Calcium imaging

A 5 mM stock solution of the Ca^2+^ indicator probe Rhod-4 AM ester (AAT Bioquest, Inc., Pleasanton, CA, no. 21120), in high-quality, anhydrous dimethyl sulfoxide (DMSO; Sigma-Aldrich, St. Louis, MO, no. D 8779) was prepared. On the day of the experiment, a concentration of 5 μM Rhod-4 AM dye in RPMI / 10% FBS media was added to the cells cultured on coverslips and incubated for 1 h. Following incubation, the media was replaced with standard RPMI / 10% FBS media without the dye. Cells were then subjected to uS stimulation as described. Cells on coverslips were then imaged using a Zeiss microscope using a Rhodamine B filter (Carl Zeiss Microscopy, LLC, White Plains, NY) for fluorescent detection and bright-field for cell visualization.

### 2.5 Biochemical polarization of macrophages

Cells were treated with 20 ng ml^−1^ IFN-γ (Sigma-Aldrich, no. I3265) and 100 ng ml^−1^ LPS (*E. coli* 0111:B4 strain, Sigma-Aldrich, St. Louis, MO, no. L2630) for 48 h for M1 polarization. and IL-4 (Sigma-Aldrich, St. Louis, MO, no. I4269) and IL-13 (Sigma Aldrich, St. Louis, MO, no. I1771) concentrations 50 ng ml^−1^ for 48 h to induce M2 polarization.^[34]^ Each of these control experiments was run three times, with comparable results.

### 2.6 Immunohistochemistry

For fluorescent imaging of iNOS and arginase proteins, the cells on coverslips were fixed in 200 μl of 4% paraformaldehyde in phosphate-buffered saline (PBS; 1X, ThermoFisher, no. J61196.AP) at room temperature for 30 minutes, and then washed twice with PBS. Cells were then permeabilized using buffer 1% Triton X-100 (ThermoFisher, no. HFH10) for 1 h, and washed 2x with PBS. Cells were then blocked, using a 5% bovine serum albumin (BSA) blocking buffer (1 mg/ml, ThermoFisher, no. J60205.AD) for 1 h at room temperature, and washed 2x with 1x PBS. Then, a primary rabbit polyclonal antibody against mouse iNOS (Cell Signaling Technology, Danvers MA, no. 68186) or mouse arginase (Cell Signaling Technology, no. 93668), a marker for M1, was incubated overnight in PBS (1:500 dilution) at 4 °C. The following morning, a secondary goat anti-rabbit IgG conjugated to Alexa-Fluor 647 (Cell Signaling Technology, no. 4414) in PBS at 1:1000 dilution was added for 2 h at room temperature in the dark. The cells were then washed 3 times in PBS. A 5 μg ml^−1^ solution of 4′,6-diamidino-2-phenylindole (DAPI; Cell Signaling Technology, no. 4083) was then added to the cells for 10 minutes. The DAPI stain was removed and cells were washed 3 times with 200 μl of 1x PBS. 10 microliters of VectaShield solution (antifade mounting medium, Vector Laboratories, Inc., Newark, CA, no. H-1000-10) was placed on a glass microscope slide and the treated coverslip was placed cell-side down onto the glass slide prior to imaging. The aforementioned iNOS experiments were run approximately *n* = 25 times (*i.e.* on *n* independent cell cultures), while the arginase controls were run *n* = 3 times.

For streptavidin-fluorescein isothiocyanate (SA-FITC) labeling of PEG-biotin-coated pzNPs, the cells were blocked using BSA and then a 0.5 μg ml^−1^ SA-FITC (ThermoFisher, Inc., no. SA10002) solution in PBS was applied to pzNP-loaded or -unloaded cells at 4 °C overnight. For simultaneous labeling of iNOS and SA-FITC, both reagents were mixed together prior to labeling. The cells were then subsequently washed in PBS prior to imaging.

### 2.7 Fluorescent imaging

For fluorescent imaging, the Zeiss Axio-Imager Z2 microscope with an Excelitas Technologies X-Cite LED 120 light box was used. The microscope was fitted with ApoTome.2 optical sections, model MCU 2008 stage controller, and model 232 power supply, all from Zeiss. For identifying M1 polarization of macrophages, a Texas Red fluorescent filter was applied. For calcium imaging of samples, a Rhodamine B fluorescent filter was applied. The images were taken with a 63x objective under oil immersion. All images within a particular experiment were acquired using identical settings (*e.g.*, laser power, gain, exposure time) on the Zeiss Z2 microscope and ZEN software. Fluorescent images were acquired using phase DIC, DAPI and fluorescent filters. Z-stack image acquisition was used with ZEN software to obtain images with 2 μm *z*-increments.

### 2.8 Statistical analysis

The mean fluorescence intensity of cells within a region of interest ROI was evaluated and quantified using FIJI/ImageJ software. For statistical comparisons, the normalities of the data distributions of the imaged data were analyzed by histogram plot analysis and the Shapiro-Wilk test (samples sizes were too small for use of the D’Agostino-Pearson or Anderson-Darling tests). This was followed by ANOVA analysis (using GraphPad Prism v.10.6.0(890) software) to determine respective *p*-values. *P*-values changed by less than 10% between ANOVA tests with and without *post hoc* analysis. Data are plotted as average fluorescence intensity within *n*

= 3 to 4 ROIs, with 20-100 cells per ROI, along with respective intensity standard deviations. Statistical significance levels are indicated directly on the figures, using the following convention: *p* < 0.05:*, *p* < 0.01:** and *p* < 0.0001:***. A *t*-test (95% confidence level) was used to compare the (IFN-γ + LPS) and (+NP, +uS) data in Figure 4, with a statistical result of *NS*: no significant difference (*p*>0.1).

## 3. Results

### 3.1 Fluorescent pzNP uptake into M0 state macrophages

An aliquot of purified, sterile pzNPs was added to RAW264.7 cells on coverslips and cultured for 48 h to allow for cellular uptake. We then performed additional image analysis for the senescent cells, including *n* = 3 replicated confocal *z*-stack slice reconstructions and images showing pzNPs in the cytoplasm of cells. **Figure 2** shows four images in different focal planes (in 2 μm height increments) of the same cells, allowing us to identify the presence and location of NPs within the cells. The results show that single or small clusters of NPs can be observed localized with cultured RAW264.7 cells sitting on the bottom of the cell culture well. Upon different focal plane sampling of the same preparation, we observe NPs internalized within the cytoplasm of the cells as well as potentially on the surfaces of the cells. The results demonstrate that, under these experimental conditions, the majority of our cells contained internalized, declumped NPs. That is, of the nine cells in the unbiased sample shown in Figure 2, all possess NPs, and this high percentage condition is representative of each of the high magnification (63×) observations made for numerous (*n*>50) NP@cell preparations. Manual counting yielded 5 - 15 pzNP/cell. The NP preparations exhibited minimal toxicity or cell death. That is, we determined that uS power density of 0.1 W cm^−2^ was sufficient to induce calcium influx and M1 polarization (iNOS expression) of RAW264.7 cells while not inducing significant cell death. We note that this ultrasound power density is comparable to that reported in the literature for pzNP excitation.^[36,44,48,53]^ We determined, via manual count of cells from a number of replicates, each before and after application of uS stimulation, that approximately 3/4^ths^ of the starting population of cells remained viable after uS, meaning they remained attached to the plates/coverslips and were available for NP and calcium uptake, cell division and polarization. The experimental conditions of applying sonication in PEG-biotin salt solution to pzNPs followed by purification of single pzNPs and a 48 h incubation was compatible with efficient cellular uptake across the majority of cells in culture.

**Figure 2.**
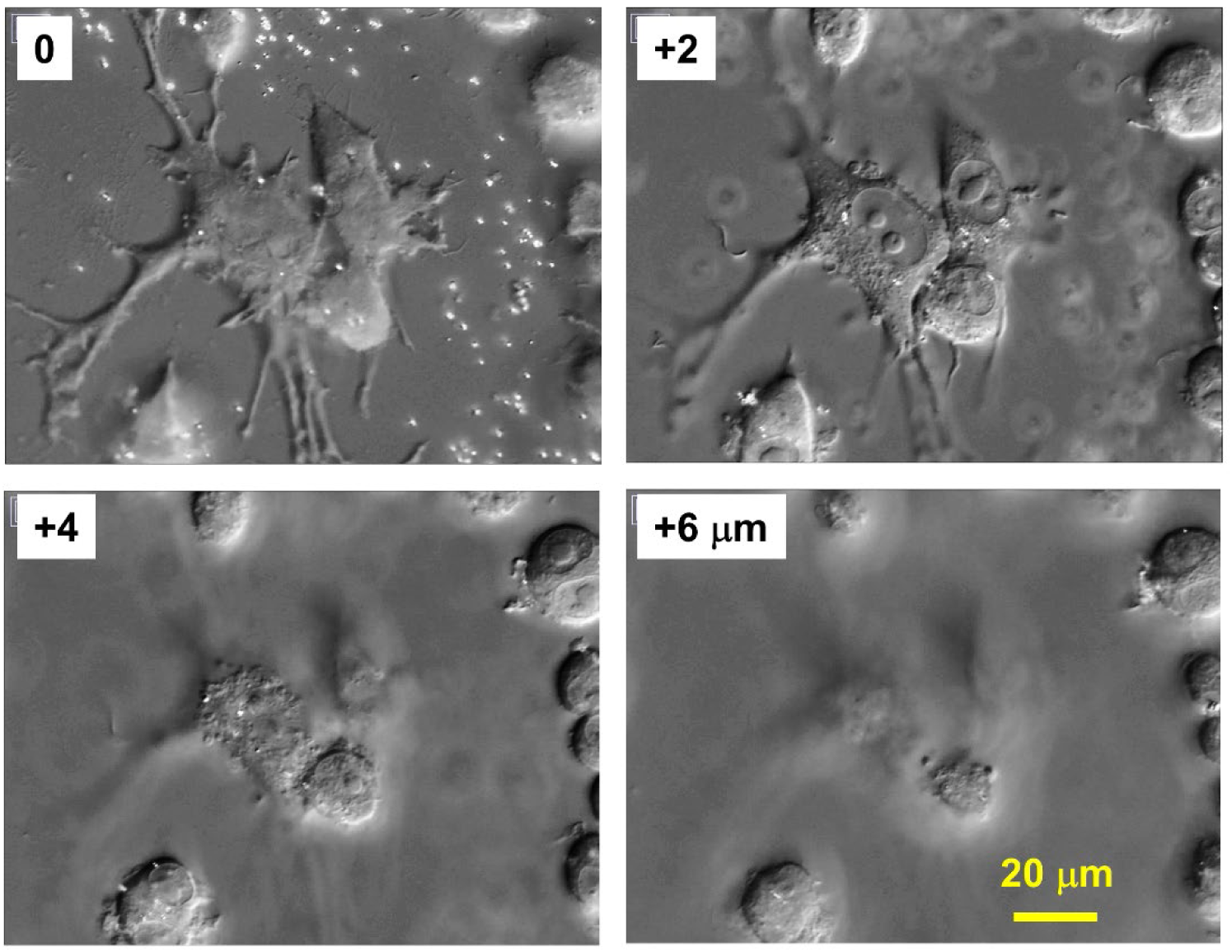
*Z*-stack confocal microscope images of RAW264.7 cells incubated with pzNPs for 48 h in media, after exposure to uS, demonstrating NP uptake by cells. Focal plane distances are shown, relative to the coverslip top surface (*i.e.* bottom of cells). Scale bar shown applies to all images.

### 3.2 Piezoelectric nanoparticle activation of calcium influx

To determine whether pzNPs, upon uptake by macrophages, can be stimulated using externally-applied uS to induce polarization, we investigated Ca^2+^ influx and cytokine expression associated with M1 activation. For calcium imaging, we used Rhod-4 AM dye that exhibits cell loading and calcium response while maintaining the spectral wavelength of Rhod-2.^[35]^ Rhod-4 AM has cellular calcium response that is 10 times more sensitive than Rhod-2 AM. The cells were incubated with pzNPs for 48 h to allow for cellular uptake. Following NP loading, the cells were incubated with Rhod-4 AM dye in RPMI media for 1 h at 37 °C to allow for uptake of the dye. The medium was exchanged and then the cells were subjected to uS stimulation. The cells on coverslips were then immediately imaged using a Zeiss Z2 Axio-Imager to monitor calcium influx.

Notably, as demonstrated in **Figure 3**, the application of uS treatment to pzNP-loaded cells (indicated as “+NP, +uS”) significantly increased the influx of Ca^2+^ relative to the (-NP, -uS) control, while pzNP-loaded cells that were not subjected to uS (+NP, -uS) and cells without NPs but with uS (-NP, +uS) exhibited no significant increase in Ca^2+^ influx (indeed, no changes with statistical uncertainties). These results demonstrate that pzNP-loaded cells exhibit a voltage-gated channel-mediated influx of calcium in response to uS stimulation. The results are consistent with pz activation of voltage-gated channels that have been previously described across multiple cell types including macrophages.^[36,37]^

**Figure 3.**
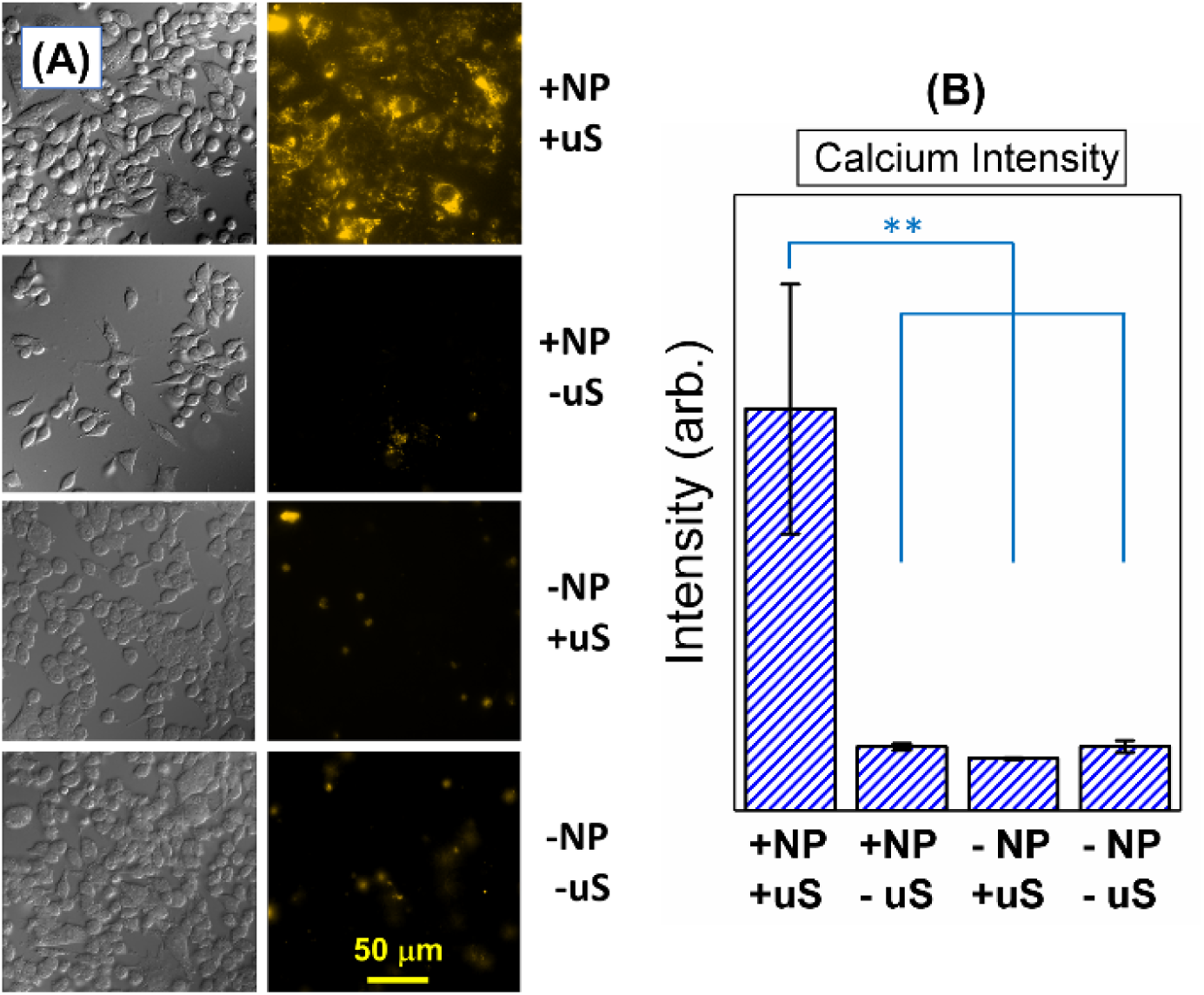
**(A)** Fluorescence images of RAW264.7 cells under various conditions, as evidenced by phase (column 1) and 4’ Rhod-4 AM calcium dye (column 2). Rows 2-4 shows a negative effect in response to (+NP, –uS), (–NP, +uS), and (–NP, –uS), respectively, whereas Row 1, (+NP, +uS), exhibits a positive effect as evidenced by a higher concentration of intracellular calcium compared to the others. **(B)** Average mean fluorescence intensity of cells within regions of interest from sets of *n=*3 fluorescent images quantified using FIJI/ImageJ, for each NP/uS condition shown. A *p*-value < 0.01 (indicated as **) in comparison to the (+NP, +uS) data was determined via ANOVA analysis. See text and Statistical Analysis for further details.

### 3.3 M1 polarization of RAW264.7 cells

As mentioned, macrophages present in different tissues are polarized according to changes in their local environment, forming different phenotypic subtypes, such as M1 and M2.^[4]^ The microbial components LPS and IFN-γ can drive macrophage polarization to the M1 phenotype while a combination of IL-4 and IL-13 can induce polarization to M2.^[4,9,33]^ M1 macrophages are capable of pro-inflammatory responses and produce pro-inflammatory related factors such as IL-6, IL-12 and tumor necrosis factor.^[1,2,38]^ In contrast, M2 macrophages elicit an anti-inflammatory response and participate in the repair of damaged tissues.^[2,39]^ In order to assess whether pzNP uptake and uS would induce a polarization phenotype of the RAW264.7 cells, we conducted a series of experiments leveraging immunohistochemistry with antibodies targeting select M1 and M2 biomarkers. The cells were plated on coverslips and then incubated with pzNP preparations for 48 h in order to facilitate efficient NP uptake prior to uS stimulation. Following a 48 h incubation with pzNPs, the samples were subjected to two rounds of uS stimulation, as per above, in order to activate the pz properties of the NPs. The cells were then cultured following uS stimulation for an additional 48 h in order to allow for efficient reproducible expression of M1 or M2 biomarkers prior to the cells being treated for subsequent immunohistochemical staining and image analysis.

The results of these experiments, presented in **Figures 4** and **5**, show that biochemical treatment of cells with IFN-γ and LPS for 48 h induced the expression of the iNOS marker of M1 polarization, as shown previously.^[1,2]^ Importantly, treatment of pzNP-loaded cells with two rounds of uS stimulation also induced the expression of iNOS, indicative of bioelectrically-driven M1 polarization (Figure 4). As shown in Figure 5, uS treatment of the pzNP-loaded cells did not induce the expression of arginase, suggesting that pzNPs do not induce M2 polarization. Also, uS treatment of the cells without pzNP preloading, and cells cultured with pzNPs but without uS stimulation, exhibited baseline-level (-NP, -uS) iNOS or arginase expression. However, as expected, control cells treated with IL-4 and IL-13 did induce arginase expression / M2 polarization, Figure 5.

**Figure 4.**
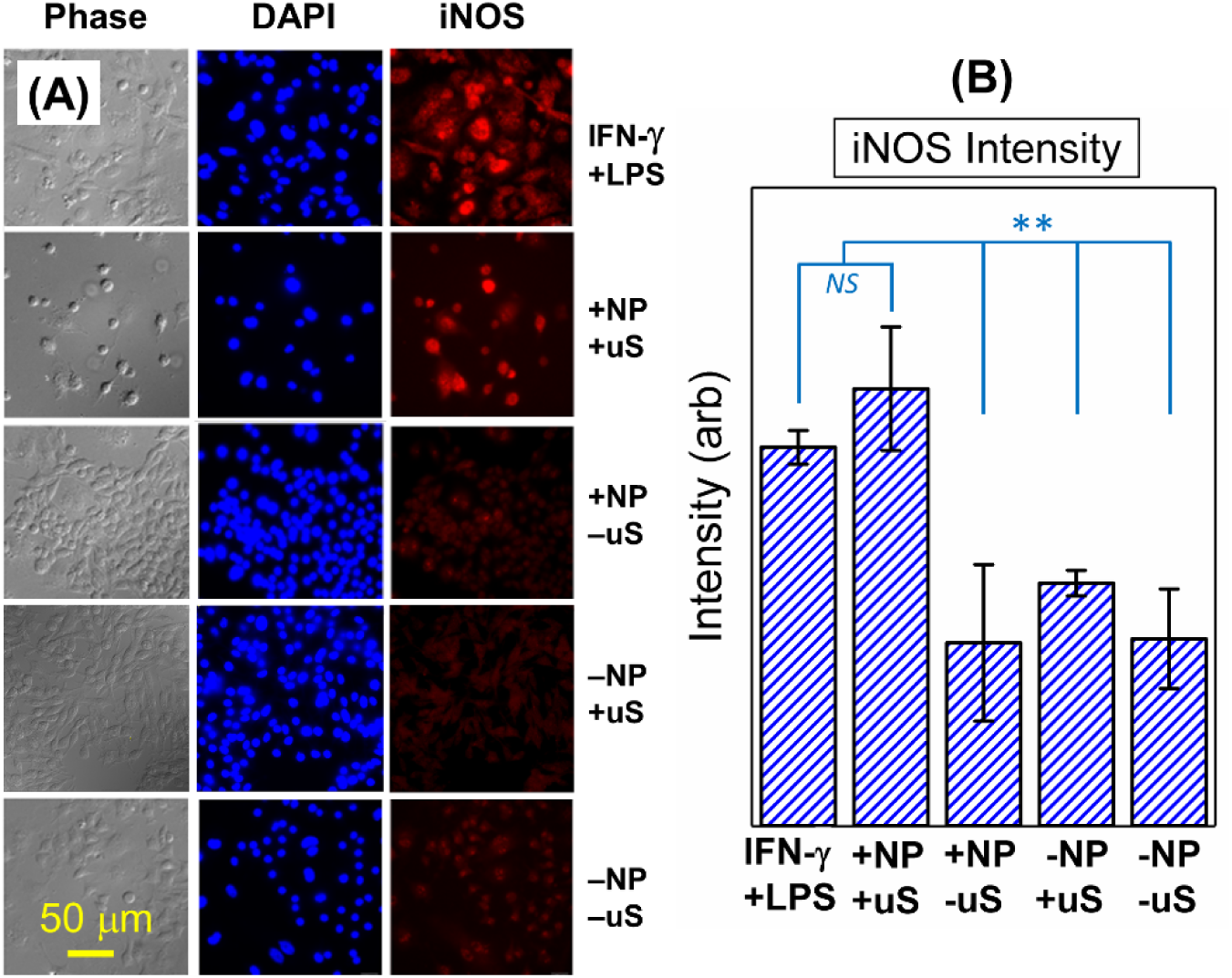
**(A)** Fluorescence images of RAW264.7 cells under different conditions demonstrating bioelectrically-driven differentiation of M0 monocytes into M1 phenotype macrophages. Phase (Column 1), DAPI staining intensity (Column 2), and iNOS staining intensity (Column 3) are shown. Row 1 shows biochemical treatment with IFN-γ + LPS that drives M0 to M1. Row 2 shows positive iNOS intensity effect with pzNP and uS (+NP, +uS). Rows 3, 4 and 5 represent controls with less / negligible iNOS staining intensity: with pzNPs and without uS (+NP, −uS, Row 3), without pzNPs and with uS (−NP, +uS, Row 4), and without pzNPs or uS (-NP, -uS, Row 5). Note that the bioelectric combination of pzNP uptake and uS inducing iNOS expression in Row 2 is similar to biochemical treatment with IFN-γ + LPS in Row 1, both indicative of M0→M1 differentiation. **(B)** Average mean iNOS fluorescence intensity of cells within regions of interest from sets of *n=*3 fluorescent images quantified using FIJI/ImageJ, for each NP/uS condition shown, plus that for IFN-γ + LPS. A *p*-value > 0.1 (indicated as *NS*, for Not Significant) was determined via a *t*-test comparing the IFN-γ + LPS and (+NP, +uS) data, while *p* < 0.01 (indicated as **) when comparing those with the three control data sets, as determined via ANOVA analysis. See text and Statistical Analysis for details.

**Figure 5.**
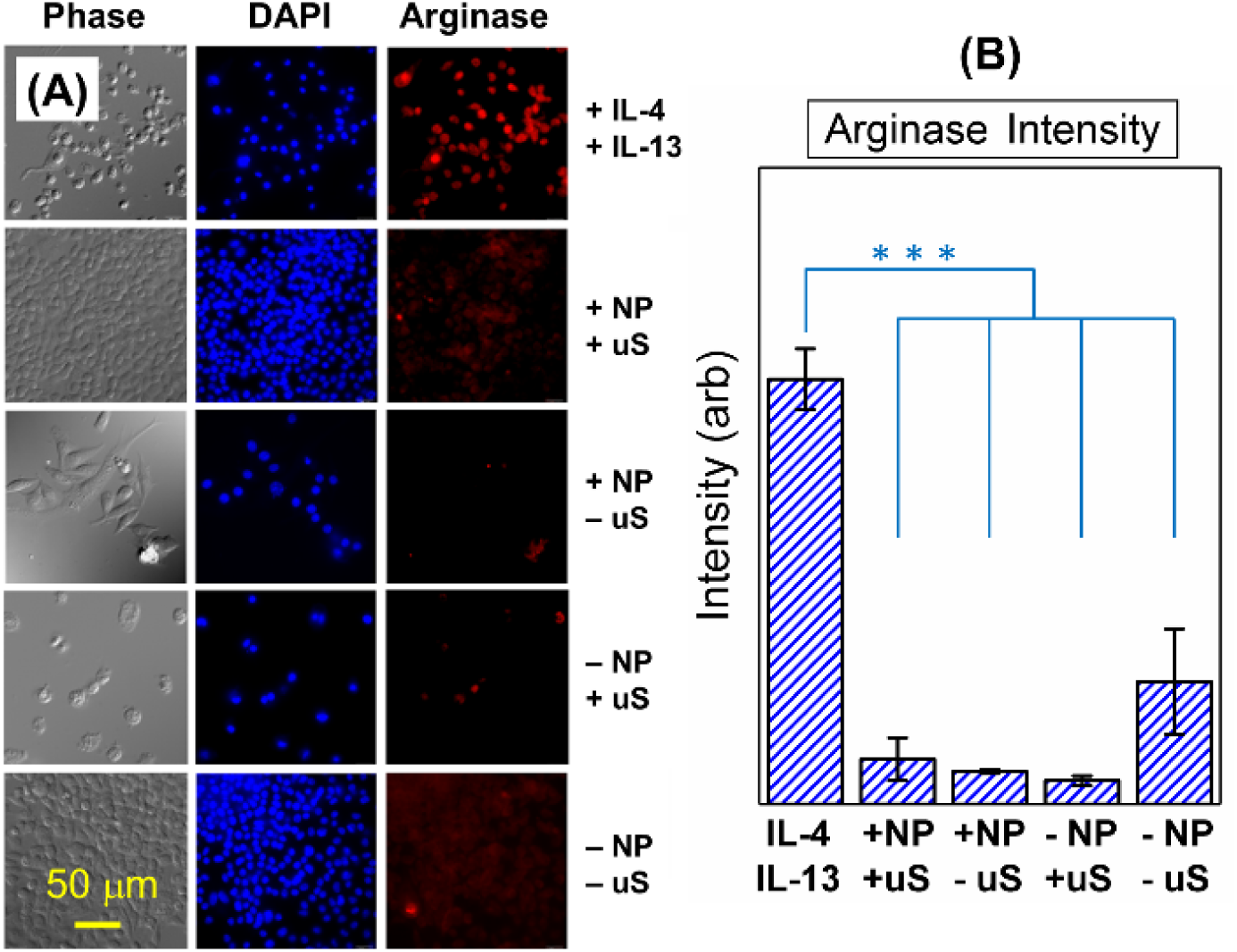
**(A)** Fluorescence images of RAW264.7 cells under different conditions demonstrating that the above bioelectric process does not induce M0→M2 polarization. Phase (Column 1), DAPI (Column 2), and Arginase staining (Column 3) are shown. Cells treated with IL-4 and IL-13 to induce M2 polarization are in Row 1, followed cells containing pzNP and uS treatment (+NP, +uS, Row 2), with the lack of red fluorescence confirming the absence of M2 polarization, and by controls with pzNPs and without uS (+NP, −uS, Row 3), without pzNPs and with uS (−NP, +uS, Row 4), and without pzNPs or uS (-NP, -uS, Row 5). **(B)** Average mean fluorescence intensity of cells within regions of interest from sets of *n=*3 fluorescent images quantified using FIJI/ImageJ, for each NP/uS condition shown, plus that for IL-4 + IL-13. A *p*-value < 0.0001 (indicated as ***) in comparison to the (IL-4 + IL-13) data was determined via ANOVA analysis. See text and Statistical Analysis for further details. See text and Statistical Analysis for details.

### 3.4 Detection of pzNP-loaded and M1-polarized cells in mixed cell populations

These results above strongly suggest that uS activation of BT pzNPs induces iNOS expression, indicative of M1 polarization of RAW264.7 cells. While this demonstrates that biophysical stimulus by piezoelectricity can modulate macrophage phenotype, the precise mechanism by which cells respond remains obscure. Toward resolving this issue, we thus attempted to leverage the immunohistochemistry assay to identify simultaneously which cells had taken up pzNPs and which cells exhibited iNOS expression. Our purification protocol entailed the use of a PEG-biotin coating of the NPs prior to cell uploading. We investigated whether we could detect uploaded NPs in cells by staining with an SA-FITC labeling protocol. We repeated the pzNP+uS stimulation experiments; however, this time, we mixed approximately equal proportions of RAW264.7 cells containing no pzNPs and cells that had been cultured with pzNPs for 48 h using coverslips. The cells adhered to the coverslips/culture dish for 24 h and were then subjected to two rounds of uS stimulation, as previously described. Following the second stimulation, the cells were fixed and stained for both iNOS expression and pzNP uploading, using SA-FITC and anti-mouse iNOS antibody. A composite fluorescence image of RAW264.7 macrophage cells following uptake of pzNPs and subsequent biophysical stimulation via uS is shown in **Figure 6**. The fluorescent images indicate DAPI blue for nuclear staining and the presence of cells, SA-FITC green for the presence of pzNPs, and iNOS red expression using an Alexa-Fluor 647 conjugated antibody. The image shows a proportion of cells containing no pzNPs and no iNOS expression (Figure 6, yellow circles) and a proportion of cells exhibiting FITC staining, indicating NP uploading (Figure 6, green circles). In addition, we detect a proportion of cells exhibiting both FITC fluorescence and iNOS expression (Figure 6, white circles). We do not detect iNOS expression in cells lacking FITC fluorescence, demonstrating that pzNP uptake to cells is a prerequisite for inducing iNOS expression. Intentionally co-culturing mixtures of cells containing NPs and without uS exposure, we can identify distinct subsets of cell populations within our *in vitro* polarization assay; namely, cells with no pzNP uptake, cells with pzNP uptake and not exhibiting iNOS expression following uS stimulation, and cells with pzNP uptake and exhibiting iNOS expression following uS. One hundred cells were analyzed for fluorescent expression, of which 50% of the cells were designated in the experiment to not have nanoparticle uptake, as discussed. In this population, we found approximately 34 stained positive for NP uptake and 17 expressed iNOS as exhibited by red fluorescence. We do not observe FITC fluorescence in the absence of NP delivery. The results further demonstrate that pzNP loading/uptake is required for the cells to induce iNOS expression upon uS simulation.

**Figure 6.**
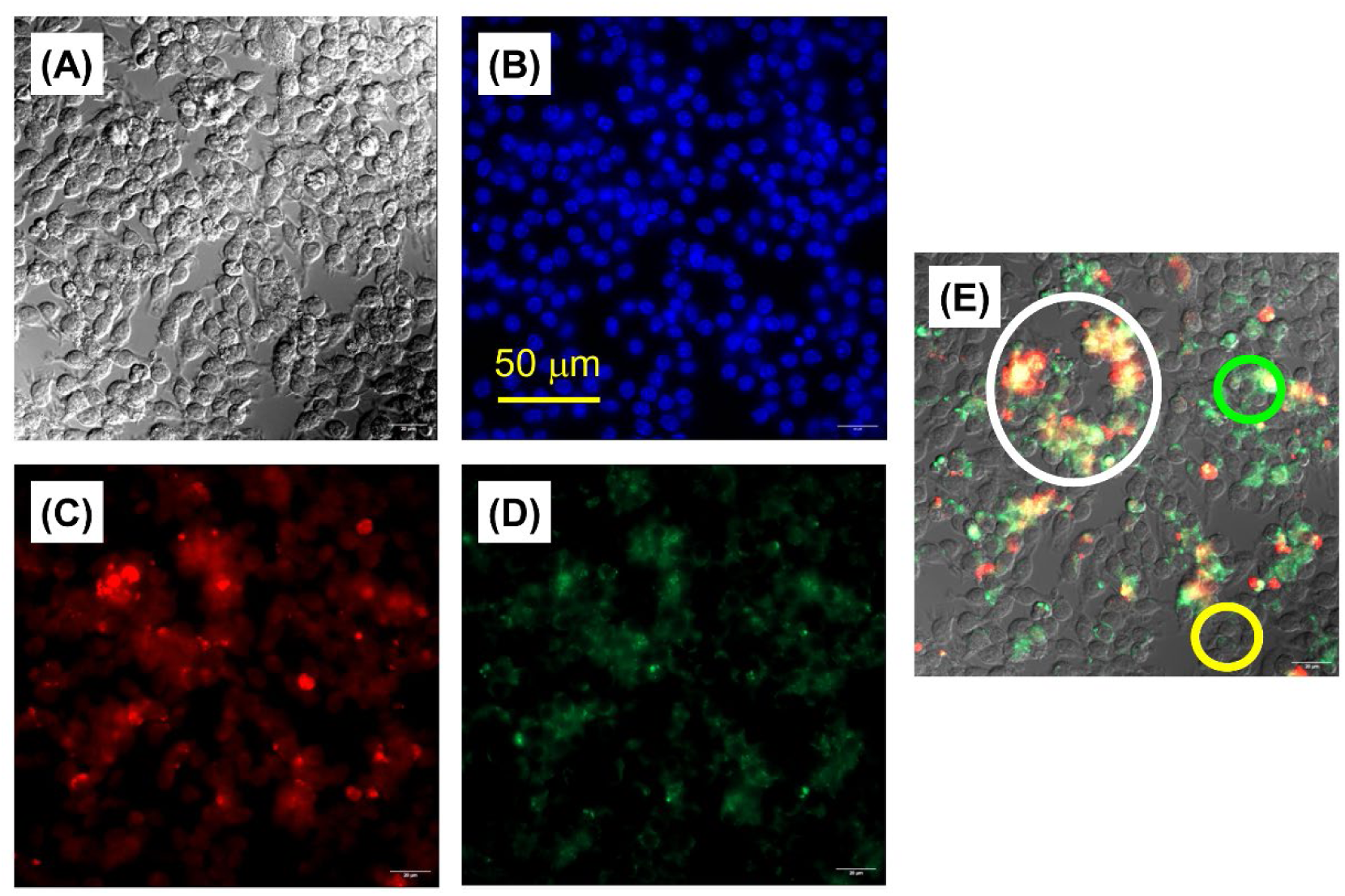
Composite fluorescence images of a mixture of RAW264.7 macrophage cells exposed and not exposed to piezoelectric nanoparticle (pzNP) uptake, and then subjected to ultrasound (uS) stimulation. The channels displayed include phase contrast (panel A), DAPI (blue, panel B) for nuclear staining, iNOS expression (red, panel C), and SA-FITC coating of pzNPs (green, panel D), in cells subjected to pzNP uptake with uS stimulation. (E) A mixture of nanoparticle and non-nanoparticle loaded cells are detected. The results indicate that discrete subpopulations of cells exposed to pzNPs and uS can be identified. Cells that have taken up pzNPs but do not exhibit iNOS expression (green circle) can be differentiated from subpopulations that have taken up pzNPs and exhibit iNOS expression (white circle). There are subpopulations of cells that were not exposed to pzNPs and when subjected to ultrasound stimulation do not express iNOS expression (yellow circle).

The data in Figure 6 further demonstrate that uS stimulation of pzNPs has the capability to induce M1 polarization *in vitro* using mouse macrophage RAW264.7 cells. We also observed that, under the conditions employed, pzNP+uS does not induce an alternative M2 anti-inflammatory cell phenotype, as measured by arginase expression (Figure 5). The M1 results demonstrate that pzNP loading of cells and subsequent uS stimulation of loaded cells immediately induces an influx of calcium ions into cells, demonstrating that the PEG-biotin-coated pzNPs are responding to uS and presumably opening voltage-gated ion channels required for macrophage polarization. In this scenario, uS waves are exploited to mechanically activate the pzNPs, thus remotely generating local electrical charges within cells by exploiting the direct piezoelectric effect. The results suggest that pz+uS stimulation and electrical charge generation within cells is activating voltage-gated ion channels consistent with the increased Ca^2+^ influx that we observed here.

## 4. Discussion

The electrical response as shown herein appears to mimic the endogenous electrical microenvironment of cells, which has also been shown capable of promoting wound healing, bone reconstruction,^[13, 40, 41]^ and nerve regeneration.^[^ ^42^ ^]^ Hence, inherent electroactive biomaterials, especially pz materials, have received wide interest as potential therapeutic modalities.^[43,44,45,46,47]^ The observed increased Ca^2+^ influx upon uS stimulation (Figure 3) suggests that electrical charge generation on pzNPs within cells activates voltage-gated ion channels. This is consistent with prior work in cells and tissues,^[14,24,48,49]^ where Ca^2+^ is a key second messenger in cell signaling and necessary for M1 polarization.^[50]^ In particular, pz material-mediated electrical potentials generated by β-PVDF membranes under uS facilitated Ca^2+^ influx through voltage-gated channels and activated the Ca^2+^-CAMK2A-NF-κB axis.^[15]^ This pathway promotes the nuclear translocation of NF-κB, leading to increased expression of inflammatory factors and enhancement of the proinflammatory macrophage response (M1 polarization). It is a crucial signaling cascade that contributes to inflammatory response and various diseases. The activation of NF-κB in this pathway can also influence the formation and function of paraspeckles, which are involved in regulating gene expression and modulating immune response.^[31]^ Paraspeckles, nuclear bodies found in eukaryotic cells, are formed through liquid-liquid phase separation (LLPS), also referred to as biomolecular condensation. This LLPS process involves the dynamic assembly of proteins and RNAs into distinct membraneless compartments. Paraspeckles are not static structures but rather undergo dynamic changes in response to changes in the cellular environment and signaling to activate macrophage polarization.^[51]^ It is possible that both biophysical and biological control of cell fate or cell polarization may intersect at the level of LLPS formation at key genomic loci.^[52]^ Biophysical stimuli such as pzNP+uS stimulation may thus act to influence calcium influx, NF-κB activation and LLPS formation within cells.

Conventional methods for applying electrical stimulation involve implanting conductive materials and supplying power through connected wires and electrodes, but these are prone to infection during prolonged use. Previous attempts mentioned above to modulate macrophage phenotype by culturing cells on pz membranes^[15,16,23]^ *in vitro* are not readily compatible with therapeutic delivery and the ability to control phenotype *in vivo*, as the membranes are not nanoscale in dimensions, and thus are not capable of intracellular uptake. Typically, these implantable materials lack spatial resolution and are most amenable to tissue repair.^[15]^ Ultrasound-driven pz neural stimulation was also performed by exploiting BT NPs in neurons.^[53]^ The pzNP-mediated uptake by individual cells and then activated by uS provides a localized signal that can modulate macrophage polarization in a non-invasive manner. Localized uS stimulation could be applied repeatedly to different regions of a tumor or to different tumors within the body. NP-loaded cells or NP preparations with subsequent cellular uptake could be repeatedly administered to achieve beneficial therapeutic results.

Currently, investigations of the effects of piezoelectric nanoparticles on cells at the single-cell level have not been as extensively explored. The ability to identify which cells have taken up nanoparticles and among this population which cells have adopted a change in cellular phenotype could, as demonstrated here, enable insights into the efficiency of NP uptake within individual cells, including variations in the number NPs taken up by cells. Analyzing cells responding or not to pzNP+uS stimulation allows researchers to identify how different cells within a population respond to pz stimulation, where the results of such stimulation can be masked when looking at aggregate cell data. By examining single cells, researchers can identify specific cell subsets that are particularly sensitive to nanoparticle exposure. The capability to differentiate cells that have adopted the M1 phenotype upon pzNP loading and uS stimulation from cells that did not adopt this M1 polarization phenotype, as well as from cells that contain pzNPs without uS exposure, can be leveraged to improve insight into the efficiency of piezoelectrical stimulation and what proportion of cells result in a distinct change in cellular phenotypes. It is currently unclear how similar are cell states achieved via pzNP+uS stimulation to those achieved via biological stimulation. It is also unclear how cells mechanistically respond to biophysical piezoelectric stimulation. Investigation of *in vivo* activation of macrophage phenotypes by remote NP stimulation requires labelling of cells with agents that can give rise to signals *in vivo,* that can be detected and measured non-invasively.

## 5. Summary and Future Perspectives

In this study, we demonstrated that piezoelectric barium titanate nanoparticles taken up by RAW264.7 macrophage cells induce a proinflammatory M1 activation upon ultrasound stimulation, as evidenced by an influx of calcium and proinflammatory iNOS expression. The cellular model can be considered a novel approach to investigating remote activation of macrophages to treat a variety of diseases including cancer and infection. The results also represent a new avenue of research with potential to reveal additional insights into how cells mechanistically respond to piezoelectric responsive materials and account for the inherent variability observed within biophysically-responsive cell populations in culture. We developed the capability to differentiate between cells that have taken up pzNPs and adopt an M1 phenotype and the subset of those cells that did not adopt an M1 phenotype, both upon uS stimulation, as well as cells that have pzNPs but were not subjected to uS. In recent years, single cell RNA sequencing, scRNA-seq, has provided significant insight into how individual cells respond to various stimuli and treatments,^[^ ^54, 55]^ and scRNA-seq across these distinct cell populations could yield additional insights. Like a kind of fingerprint, the number of different mRNA molecules per gene in a particular cell informs about the cell’s identity and the functional phenotype.

Future investigations could utilize the existing biological cell model described here to uncover insights into mechanistic understanding of how cells respond to piezoelectric stimulation. The scRNA sequencing approach could involve fluorescent sorting of nanoparticle-loaded cells or the coupling of unique DNA oligonucleotide barcodes to the aforementioned PEG-coated pzNPs. The experimental approach described here may offer a straightforward way to inform which genes are changing expression in response to both biological (IFN-γ+LPS) and bioelectric (pzNP+uS) stimulation, addressing the question “Are there differences between pzNP-induced and biologically-stimulated M1 cells and, if so, what are they?” An examination of single cell RNA expression datasets could provide valuable information in regard to the proportion of cells taking up pzNPs and the proportion of pzNP-loaded cells that respond to piezoelectric stimulation and adopt an M1 phenotype. We anticipate that, following the current work with exploration of single cell RNA sequencing across these populations, we will gain additional insights into the mechanism by which macrophage cells respond to piezoelectric stimulation. Without new insights into the biophysical mechanism, the intelligent design of more efficient piezoelectric control of macrophage cells will be hindered. There are a number of advantages in the exploitation of piezoelectric nanoparticles in immune cell regulation. Improved understanding of how different cells can respond to external stimuli such as ultrasound and convert this form of energy into physical, electrical or chemical cues to change a cellular phenotype has much interest. It is anticipated that these investigations will contribute to significantly improved understanding of how piezoelectric stimulation influences cell behavior, moving such research closer to therapeutic development.

## Author Contributions

**Timothy Connolly**: Methodology, Investigation, Formal analysis, Supervision, Writing – original draft, review and editing. **Camille Johnson**^‡^ and **Allison Chen**^‡^: Investigation, Formal analysis, Writing - review and editing. **Krzysztof Kempa**: Software, Investigation. **Dylan Hatt**: Investigation, Formal analysis. **Michael J. Naughton:** Conceptualization, Project administration, Methodology, Investigation, Formal analysis, Writing – original draft, review and editing.

## Acknowledgements

We thank Dr. Bret Judson and the Boston College Imaging Core for infrastructure and support, Mr. Jorel Padilla for image analysis support, and Dr. Adam Vanesse for assistance with thermal, DLS and uS intensity characterizations.

## Conflict of Interest Statement

The authors declare that they have no conflicts of interest.

## Data Availability Statement

The data will be available from the corresponding author on reasonable request.

## Ethical Statement

Not applicable.

## Notes

### Competing Interest Statement

The authors have declared no competing interest.

